# Three-dimensional assessments are necessary to determine the true, spatially-resolved composition of tissues

**DOI:** 10.1101/2023.12.04.569986

**Authors:** André Forjaz, Eduarda Vaz, Valentina Matos Romero, Saurabh Joshi, Alicia M. Braxton, Ann C. Jiang, Kohei Fujikura, Toby Cornish, Seung-Mo Hong, Ralph H. Hruban, Pei-Hsun Wu, Laura D. Wood, Ashley L. Kiemen, Denis Wirtz

## Abstract

Methods for spatially resolved cellular profiling using thinly cut sections have enabled in-depth quantitative tissue mapping to study inter-sample and intra-sample differences in normal human anatomy and disease onset and progression. These methods often profile extremely limited regions, which may impact the evaluation of heterogeneity due to tissue sub-sampling. Here, we applied CODA, a deep learning-based tissue mapping platform, to reconstruct the three-dimensional (3D) microanatomy of grossly normal and cancer-containing human pancreas biospecimens obtained from individuals who underwent pancreatic resection. To compare inter-and intra-sample heterogeneity, we assessed bulk and spatially resolved tissue composition in a cohort of two-dimensional (2D) whole slide images (WSIs) and a cohort of thick slabs of pancreas tissue that were digitally reconstructed in 3D from serial sections. To demonstrate the marked under sampling of 2D assessments, we simulated the number of WSIs and tissue microarrays (TMAs) necessary to represent the compositional heterogeneity of 3D data within 10% error to reveal that tens of WSIs and hundreds of TMA cores are sometimes needed. We show that spatial correlation of different pancreatic structures decay significantly within a span of microns, demonstrating that 2D histological sections may not be representative of their neighboring tissues. In sum, we demonstrate that 3D assessments are necessary to accurately assess tissue composition in normal and abnormal specimens and in order to accurately determine neoplastic content. These results emphasize the importance of intra-sample heterogeneity in tissue mapping efforts.

## INTRODUCTION

Recent developments in spatial profiling technologies have led to the construction of atlases to characterize cellular and tissue compositions, structure, and the “-omic” (genomic, epigenomic, transcriptomic, proteomic, and metabolomic) landscapes of tissues, organs, and whole organisms.^1–9^ These techniques have led to important discoveries regarding changes in cellular composition during development, aging and the progression of diseases such as cancer and cardiovascular disease. However, due to technical and financial limitations, current spatial omic methods – CODEX, IMC, Visium, DBitseq, seqFISH, MERFISH, etc. – are designed to evaluate mm^2^-sized two dimensional (2D) regions.^1,6,10–13^

For a histological section of standard, 4 µm thickness, a 1-mm^2^ core of a tissue microarray (TMA) represents a volume of tissue of just 0.004 mm^3^, while a common region size for spatial transcriptomics (6.5 x 6.5 mm^2^) corresponds to a volume of 0.2 mm^3^. These subsampled volumes represent minuscule fractions of the human organs and diseased regions that they are used to represent. More standard techniques, including whole slide images (WSIs) stained with hematoxylin and eosin (H&E) or immunohistochemistry (IHC), are often considered the gold standard of diagnostic anatomic pathology.^14,15^ These slides feature a lateral area of 2 x 5 cm^2^, corresponding to a volume of 5 mm^3^.

The implicit assumption of 2D sampling via TMAs or WSIs is that the cells within the sampled region, as well as their morphologies, densities, and neighborhoods, are representative of those of the three-dimensional (3D) organ from which they were sampled. Admittedly, accurate clinical diagnosis of a range of diseases using single 2D H&E sections (selectively from gross inspection of resected tissues) shows that generalization of findings from 2D is possible in cases of binary questions such as presence or absence of cancer cells. In clinical settings, 2D assessments have proved to be typically sufficient. However, in research settings, where the goal of tissue atlas efforts is generalizability, we hypothesize that 2D sampling may be insufficient to capture the marked intra-sample heterogeneity in cellular composition and tissue architecture.

Recent 3D work has demonstrated the utility of tissue clearing and serial sectioning-based approaches to assess microanatomical maps of large (>1 cm^3^) volumes of tissue at cellular resolution. ^16–30^ Here, we use the recently developed 3D imaging workflow CODA to assess the spatial composition of key cells types in thick slabs of both grossly normal human pancreas tissue and human pancreas tissue containing pancreatic ductal adenocarcinoma (PDAC), a deadly and common pancreatic neoplasm.^18^ The uniquely heterogeneous spatial microenvironment of PDAC makes it an optimal testbed to evaluate the benefits of 3D microanatomic mapping over standard 2D approaches.^31–33^

Our analyses demonstrates that standard 2D sampling – using a limited number of TMA cores or WSIs – is typically insufficient for accurate assessment of tissue composition, tumor content, or the selection of regions of interest for creation of TMA cores and capturing rare events such as separate small cluster of cancer cells in pancreatic tumors.^34^ We determine that tens of WSIs and hundreds of TMA cores are be necessary to accurately represent the range of tissue compositions present in a cm^3^-sized human pancreas sample. We find that sections inside a tumor, sometimes just tens of microns from each other, can have completely different, uncorrelated cellular and non-cellular structures. Two-dimensional assessments of “representative” slides fail particularly in enumeration of rare events, such as estimation of the density of cancer or cancer precursor cells in samples known to have low neoplastic content.^19,34^ This work aims to clarify the impact of tissue subsampling in study of the composition of normal and malignant tissues.

## METHODS

### Pancreas specimen acquisition

This retrospective study was approved by the Johns Hopkins University Institutional Review Board (IRB). Three cohorts of pancreas tissue were assembled here, which we term TMA, 2D-WSI, and 3D-CODA. The TMA was purchased from TissueArray. It contained 60 1.5 mm diameter cores taken from the pancreata of 30 individuals diagnosed with pancreatic cancer. The 2D WSI cohort contained tissue from the pancreata of 64 individuals who underwent surgical resection for pancreatic cancer at the Johns Hopkins Hospital, and is a cohort that was previously published.^35^ The 3D pancreas cohort contained tissue from the pancreata of seven individuals who underwent surgical resection for pancreatic cancer at the Johns Hopkins Hospital, and is a cohort that was previously published. ^17,18^ An additional seven specimens of normal adjacent surgically resected human pancreas tissue from individuals who underwent pancreatic resection for pancreatic abnormalities were also included.

### Tissue processing

Resected tissues were formalin-fixed, paraffin embedded, and sectioned at a thickness of 4 µm. For the TMA and 2D-WSI cohorts, a single histological section was stained with H&E. For the 3D-CODA cohort, a minimum of 270 serial sections were taken, and every third section was stained with H&E, for a minimum of 90 H&E-stained tissue sections per sample. H&E-stained images were digitized at 20x magnification using a Hamamatsu S360 scanner.

### Segmentation of pancreatic microanatomy in 2D

A previously developed deep learning semantic segmentation pipeline for the labelling of distinct microanatomical components in histological images was adapted here to label ten microanatomical components of human pancreatic cancer histology at 1 µm per pixel resolution: pancreatic cancer, pancreatic cancer precursor lesions, normal ductal epithelium, acinar tissue, islets of Langerhans, vasculature, nerves, fat, and stroma.^18,36^ Convolutional neural networks were trained in MATLAB2023b to classify the TMA cores, 2D-WSI, and 3D-CODA cohorts of tissues. Manual annotations of the ten microanatomical tissue components were generated on a subset of histological images, and fed into the CODA-segmentation workflow for retraining of a resnet-50 network. Resulting networks were deemed acceptable if the overall accuracy exceeded 90% and minimum per-class precision and recall exceeded 85%.

### Reconstruction of pancreatic microanatomy in 3D

CODA image registration was used to create digital tissue volumes from the serial H&E images for the seven samples in the 3D-CODA cohort.^18^ This nonlinear registration workflow iteratively aligns serial stacks of images (with the reference coordinates at the center of the stack), and utilizes a two-step global and local calculation in MATLAB2023b. Images are downsampled to a resolution of eight µm per pixel, converted to greyscale, and Gaussian-filtered. Global registration angle is calculated through maximization of the cross correlation of radon-transforms of the filtered images taken at discrete angles from 0 - 360°, and registration translation is calculated through maximization of the cross correlation of the rotated, filtered images. Local registration is computed by repeating this process along subsampled regions of the two globally registered images. This registration is repeated for all images in the serial samples and is subsequently rescaled and applied to the high resolution (1 µm per pixel resolution) H&E and microanatomically segmented H&E images.

### Calculation of variation in tissue composition in 2D and 3D

For each discrete sample in the TMA, 2D-WSI and 3D-CODA cohort, overall microanatomical composition was assessed. First, the number of pixels classified as each of the 10 microanatomical tissue types segmented by the deep learning model was determined. Next, composition was defined as the area percent of each tissue type in each sample. Variation in tissue composition in the 2D-WSI cohort was defined as the distribution of composition of each tissue type segmented by the deep learning model. Variation in tissue composition in the 3D-CODA cohort was defined as the distribution of composition of each tissue type along the z-dimension of the serial stack of images, taking each serial histological image as an independent measurement. Minimum, maximum, mean, median, standard deviation, and histogram bin counts of each tissue component composition in the 2D-WSI and 3D-CODA cohorts were determined. In determination of the distribution of tissue composition in the 3D-CODA cohort, samples containing >90 serial images were randomly subsampled to contain 90 consecutive images.

### Calculation of the number of tissue microarrays necessary to understand WSI and 3D tissue composition

Virtual TMAs (vTMAs) were generated in the 2D-WSI and 3D-CODA samples. First, a 2D or 3D coordinate was generated. Pixels were extracted corresponding to a 1 x 1 mm^2^ square surrounding the coordinates. A circular filter was applied to this extracted square to leave a 1-mm diameter disk representing a vTMA taken from the 2D or 3D image. To determine the number of vTMAs necessary to accurately estimate the tissue composition of a WSI or 3D pancreatic cancer tissue sample, random coordinates were determined, virtual TMAs were generated, and the tissue composition of each vTMA was recorded. Error was calculated between the per-class vTMA tissue composition and the overall composition of the WSI or 3D tissue sample. Another random vTMA was generated, added to the first vTMA, and error was recalculated for the combined sampling of two vTMAs. This process was repeated for sampling of up to 800 vTMAs on the 3D tissue samples and up to 100 vTMAs on 2D-WSIs. One thousand such simulations were performed to determine the general trend of per-class TMA error in assessment of WSI and 3D sample tissue composition.

### Calculation of the decay in spatial correlation within 2D and 3D samples

In each sample of the 3D-CODA cohort, 2D planes of pixels were extracted from each classified tissue volume. For each segmented tissue component, the cross-correlation of the pixels classified as that component in that plane to all other planes of the 3D-sample was determined, and this correlation along with the distance between the planes was recorded. This process was repeated for all possible combinations of z-planes in a volume, and was repeated for each of the seven tissue samples. Aggregate correlation of composition of a single tissue component as a function of distance within a 3D sample was defined as the mean cross-correlation of that tissue component across all images of all samples.

### Calculation of the change in tissue composition along vTMA cores

Change in tissue composition along serial sections of a vTMA core was determined. First, manual selection of coordinates on the first image of a sample was selected corresponding to a region visibly seen to contain invasive cancer. Next, a virtual core was extracted from the 3D segmented tissue volume corresponding to a cylinder of 3 mm diameter. Serial vTMAs were taken from each core, and the tissue composition of each serial vTMA was determined. Error in composition of each tissue type between the initial, manually selected vTMA and each serial TMA was calculated, and recorded along with the section number of that virtual serial TMA.

### Calculation of the number of sections necessary to understand neoplastic content

Pancreatic neoplastic content was defined in two ways. For pancreatic cancer precursor lesions, PanIN content was defined as the volume of PanIN normalized by the combined volume of PanIN and normal ductal epithelium. For pancreatic cancer, PDAC content was defined as the volume of PDAC normalized by all PDAC, epithelial ducts, and PanIN total volume of the 3D sample. For each 3D sample, subvolumes were extracted corresponding to all combinations of between 1 and 90 serial tissue images. For each unique combination, the neoplastic content of the subvolume was calculated, and the relative error of this content was determined in relation to the neoplastic content of the whole 3D volume. For each 3D sample, measurements were grouped by the number of serial images contained in each subvolume.

### Statistical considerations

All significance tests were performed using the Wilcoxon rank sum test. To compare metrics within and between cohorts, median, mean, standard deviation, and interquartile range were determined. Relative error was defined as [measured value – expected value] / expected value. No other statistical calculations were performed in this work.

## RESULTS

### Construction of cohorts of microanatomically labelled pancreatic tumors

To interrogate deeply the differences between inter-sample and intra-sampled compositional heterogeneity, pancreatic tissue from a total of 149 individuals was analyzed in this work. Pancreatic tissues containing invasive pancreatic cancer were analyzed from a total of 101 patients (Fig 1A). An additional 48 samples of grossly normal pancreas were analyzed in the non-diseased cohort. Cohorts consisted of individual, pathologist-curated WSIs containing pancreatic cancer from 64 patients (“2D-WSI” cohort, Fig 1B), and serially sectioned histological images of tumor blocks containing pancreatic cancer from seven patients (“3D-CODA” cohort, Fig 1C). A single TMA containing invasive pancreatic cancer from 30 individuals was also assessed. As discussed below, Fig. S2 indicates that the tissue composition combinations for these 64 WSIs in the 2D cohort saturates at 40-50, such that additional patients do not provide further information regarding compositional heterogeneity (note we do not consider genomic or transcriptional heterogeneity here).

**Fig. 1.**
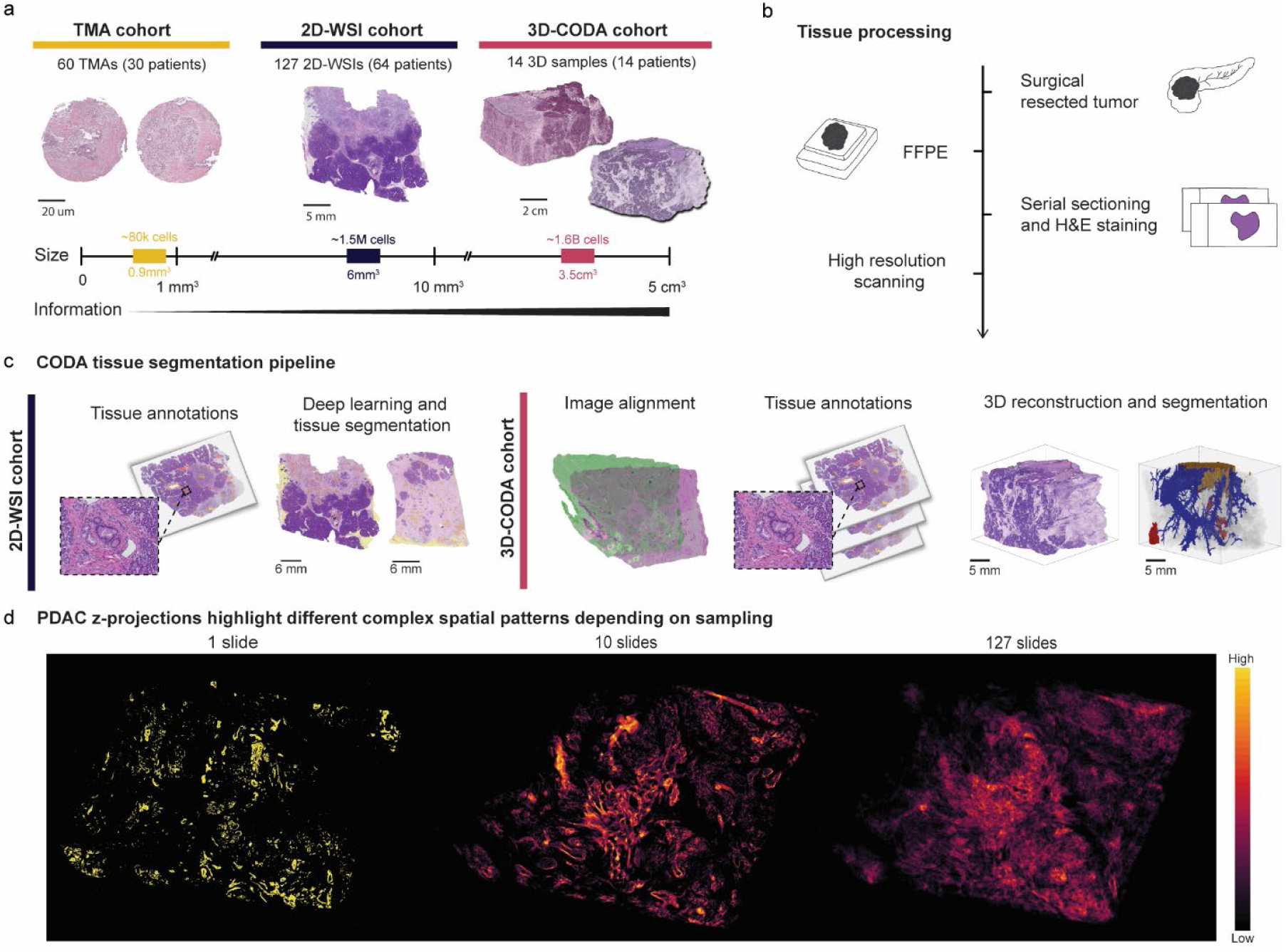
Construction of cohorts of microanatomically labelled pancreatic cancer histological sections for assessment of inter-and intra-patient tumor heterogeneity. **(a)** Cohorts of 3D blocks, 2D-WSIs, and 2D-TMAs from surgically resected human pancreatic tissues were retrospectively collected from 101 individuals diagnosed with pancreatic cancer. A cohort of 127 specimens from 64 different patients were analyzed as single whole slide (“2D-WSI” cohort; 1-2 slides per patient). A cohort of 14 blocks were serially sectioned and reconstructed in 3D using the AI-based platform CODA (“3D-CODA” cohort). A cohort of 60 TMA cores were serially sectioned and segmented using CODA (TMA cohort).**(b)** All tissue cohorts were surgically resected, histologically sectioned, stained with H&E and digitized.**(c)** For the processing of the 2D-WSI cohort, CODA segmentation was used to label 10 different microanatomical components (including epithelial ducts, fat, islets of Langerhans, PDAC, acini, nerves, blood vessels, and extracellular matrix (ECM)) at a resolution of 1 micron. For the processing of the 3D-CODA cohort, serially sectioned specimens were registered into aligned tissue volumes. Manual annotations of a subset of images were used to train a deep learning model to automatically segment anatomical labels and subsequently reconstructed them in 3D. **(d)** Local PDAC content projected in 2D from a stack of 127 sections shown as a heatmap to exemplify the spatial distribution of cancer in 3D samples. Selection of single whole slide from the stack of WSIs (left), selection of ten whole slides (middle), and all whole slides in the sample (right). Higher PDAC content on the z-axis is highlighted in yellow regions and low PDAC content is labelled in black regions.

For each cohort, a trained semantic segmentation algorithm was used to label microanatomical components to a lateral resolution of 1 µm. Independent assessment of the trained model revealed an overall accuracy of 93.2% (Fig S1). For the 3D-CODA cohort, image registration was performed on images of serially cut sections to create 3D digital tissue volumes (Fig 1B). The minimum number of serial sections per sample was 270 sections (mean: 297 sections, interquartile range: 816 sections). The median reconstructed sample volume was 39.0 mm^3^ (mean: 132.2 mm^3^, interquartile range: 247.3 mm^3^).

Fig 1D displays z-projections of PDAC highlighting different complex spatial patterns depending on sampling. The heatmaps were generated by segmentation of an aligned serial sectioned stack of 127 WSIs. XY locations with higher PDAC content on the Z (axial) dimension are highlighted, showing greater spatial heterogeneity across planes. The heatmap with one single whole-slide (left) shows small sparse sampling of PDAC, the heatmap for 10 serial sections (middle) of the tissue block reveals additional PDAC spatial patterns, while the heatmap containing the full tissue block (right) provides a more comprehensive representation of the PDAC spatial tissue heterogeneity.

### Comparison of intra-tumoral and inter-tumoral heterogeneity in microanatomical composition

To compare the heterogeneity in tissue composition present within the 2D and 3D cohorts, we quantified the relationship between inter-tumoral differences in tissue content, as measured in the 2D-WSI cohort, and intra-tumoral differences in tissue content, as measured in the 3D-CODA cohort. First, we determined the per-tissue component composition of each image in the 2D-WSI and 3D-CODA cohorts using CODA. The tissue components segmented by CODA in the 2D-WSI and 3D-CODA cohorts included PDAC, non-neoplastic ductal epithelium, islet of Langerhans, blood vessel, extracellular matrix (ECM), acini, fat, and nerve. To minimize sample-size bias, any 3D-CODA sample that contained >64 serial sections was randomly subsampled to 64 consecutive sections. Next, we analyzed the range in tissue content for each of the tested tissue component in the 64 samples of 2D-WSI cohort (range across all 64 samples in black, Fig. 2A) and for each of the samples where that component was present in the 3D-CODA cohort (range for each sample in red, Fig 2A). To account for the arbitrariness of our sampling of the seven 3D samples of pancreatic cancer, we subsampled the 3D cohorts to all possible combinations of one through seven samples.

**Fig. 2.**
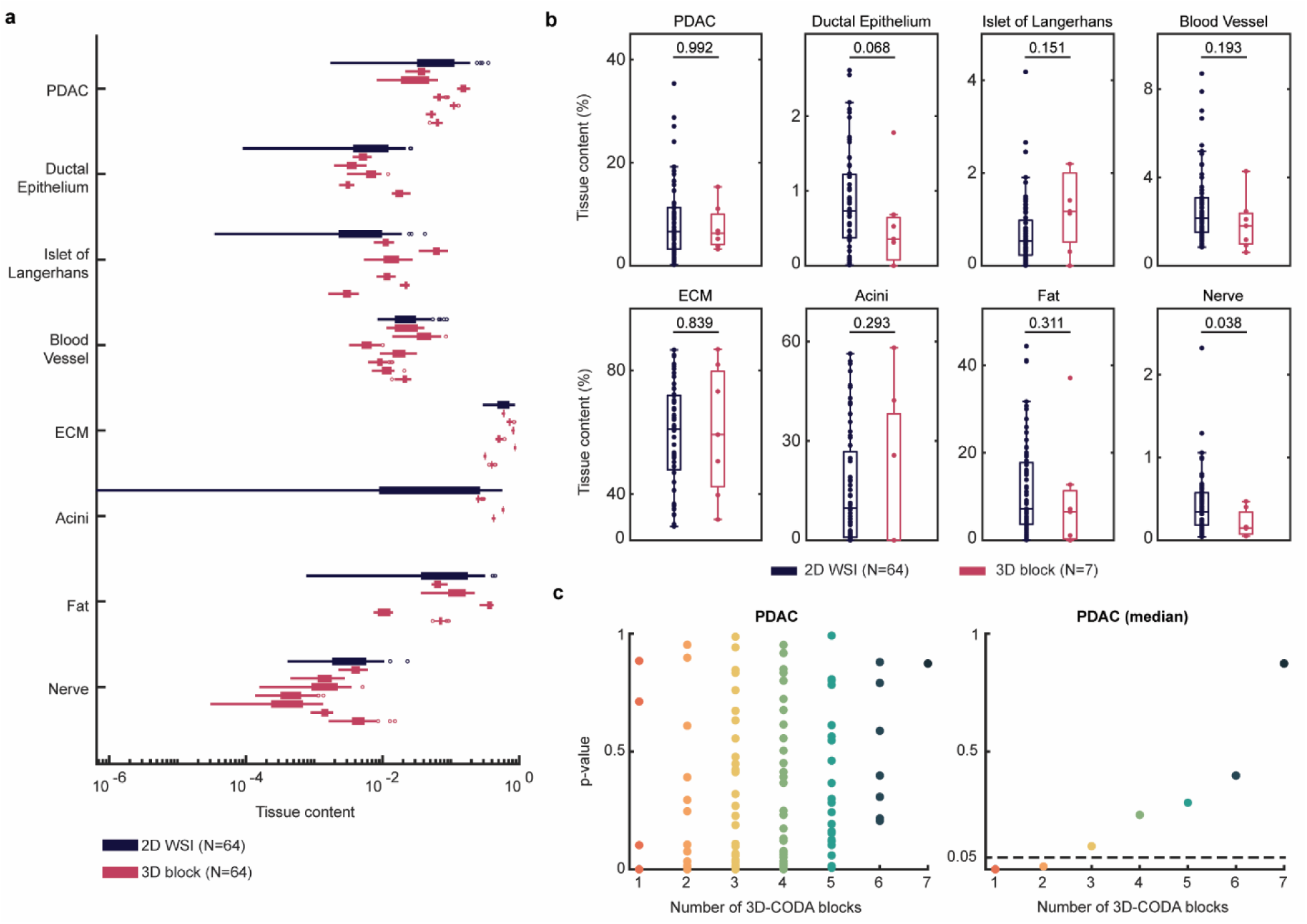
Comparison between inter-and intra-patient tumoral heterogeneity in tissue composition. (**a**) Range in bulk tissue content for 64 slides from the 2D-WSI cohort (black) and the 7 samples (each containing 64 slides) of the 3D-CODA cohort (red). The tissue components segmented and analyzed by CODA for both cohorts include PDAC, ductal epithelium, islet of Langerhans, blood vessel, extracellular matrix (ECM), acini, fat, and nerve. Note that tissue components are not all present in all 7 blocks, which is why the number of blocks shown is different for different tissue components. (**b**) Comparison between the percent of each tissue component for each section of the 2D-WSI cohort and for the blocks of the 3D-CODA cohort. This reveals non-significant differences in inter-and intra-patient tumoral heterogeneity between the two cohorts across all tested tissue components, besides nerve. (**c**) 2D-WSI tissue composition range is compared to 3D-CODA composition range for all possible combinations of one through seven 3D samples (left panel), revealing that the median difference between the 2D and 3D cohorts become non-significant (p>0.05) with as few as three 3D samples (right panel).

These data indicated that the wide distribution in tissue content present across 64 independent WSIs (one per patient) was nearly fully represented by the heterogeneity of the seven 3D samples. Besides nerve, the inter-patient heterogeneity in each tissue component of the 2D-WSI cohort was statistically insignificant when compared to the intra-patient heterogeneity of the seven 3D-CODA samples (Fig 2B).

Remarkably, this analysis revealed that the range in heterogeneity observed in just three 3D samples was sufficient to replicate the range of inter-patient heterogeneity in tissue composition (exceed a p-value of 0.05) within the 2D cohort (Fig 2C). The occurrence of PDAC tissues present in the 2D-WSI cohort was compared to that of all combinations of samples present in the 3D-CODA cohort to assess statistical significance of the overlap between the two cohorts (Fig S2).

Finally, to validate the number of WSIs we chose in the 2D-WSI cohort, we calculated the heterogeneity in tissue composition for various numbers of WSIs. This analysis measured the number of 2D WSIs necessary to saturate in inter-patient compositional heterogeneity. We found that the range in content for all tissue components saturates at a number of slides between 40-50 (Fig S2B), justifying our use of 64 WSIs in this work.

### Spatial correlation in tissue composition rapidly decays within pancreatic tumors

To assess the degree of heterogeneity in tissue content within 3D samples, we calculated how much spatially resolved tissue composition changed when traveling in a straight line through a 3D tumor. To determine the correlation length of each pancreatic tissue component (PDAC, fat, etc.) – i.e. the persistence distance over which the tissue composition remained significantly correlated – we calculated the spatial (pixel-to-pixel) correlation of structures across the samples of the 3D CODA cohort (Fig 3A). This correlation was calculated for each tissue component and for all whole-slide images spaced between 4 µm and 720 µm apart in the 3D samples. Correlation was averaged across the seven 3D samples and plotted as a function of distance between slides (Fig 3B). This assessment was repeated for the eight main tissue components present in the pancreatic tumors (Fig 3B).

**Fig. 3.**
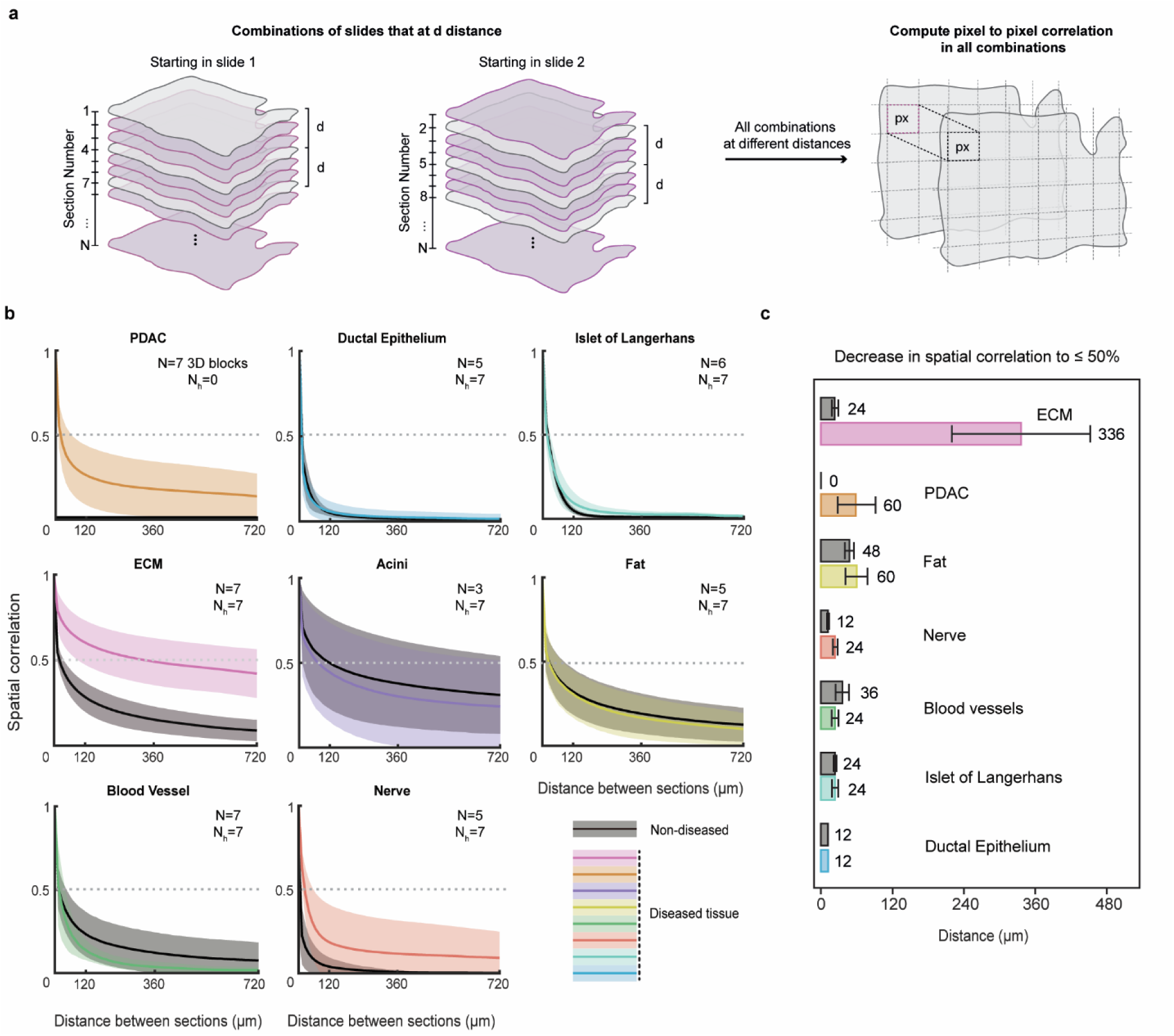
Rapid decay of the spatial correlation in tissue content within tumors. **(a)** 2D spatial correlation in tissue content was calculated for all combinations of pairs of sections within the samples of the 3D-CODA cohort. **(b)** For each tissue type, the correlation was plotted as a function of the distance between the section pairs. Spatial correlation decay was computed for diseased and non-diseased samples. Dotted line shows a correlation of 50%. Lines are the mean; shaded areas represent the standard deviations. As tissue components are not all present in all 7 blocks, the number N of blocks shown is different for different tissue components. **(c)** Distances at which the correlation falls below 50% for each of the test tissue components and each of the 14 samples in the 3D-CODA cohort (diseased and non-diseased). This shows the tissue composition in a given section in a block becomes essentially uncorrelated with tissue composition in a second slide in the same block if that slide is just tens of microns from the first one. In other words, the prevalence of tissue composition in a PDAC tumor is extremely short.

This analysis determined the extent to which each tissue maintains anatomical continuity. Our analysis of each individual tissue revealed that the more abundant tissue structures in the pancreatic tumor tissues, such as ECM and acini, remained spatially correlated over large distances within the blocks, requiring >180 slides (or 720 µm) until they reached a spatial correlation that had decreased by >50%. For sparser tissues, such as nerve and blood vessel, no more than a distance of 24 µm corresponding to just six 4-μm-thick slides, were required for the spatial correlation in the content in that tissue component to fall below <50% (Fig 3C). Correlation in ductal epithelium content dropped by 50% within 12 μm. These results reveal the significant changes that occur in tissue content across short distances.

Similar analysis of spatial correlation decay was conducted for non-diseased pancreatic specimens. The pattern of ECM heterogeneity was lower when compared to the diseased tissues, with a steep decrease of 50% in spatial correlation in just 24 μm. In contrast, acini decayed more slowly in normal pancreas tissues. This inverse pattern reflects distinct microanatomical differences between normal and cancer-containing pancreas, in particular acinar loss, and ECM deposition characteristic of PDAC progression.

A corollary of this is that sampling a 3D tumor at a rate corresponding to a distance between sections that is larger than the correlation length is insufficient for a rigorous assessment of the spatially resolved composition of that tumor. This provides guidelines for the minimum spatial rate (maximum distance between whole slides) at which the content in a specific tissue component needs to be assessed.

### Limitations of core-needle biopsies in assessment of tumor heterogeneity in tissue composition

To further quantify the loss in spatial correlation of cancer across thick slabs of tissue, we created virtual cores within the large 3D samples to simulate TMAs (Fig 4A). In specimens of the 3D-CODA cohort, 50 locations were manually chosen on the first H&E image as regions containing visually high neoplastic content. This is to mimic how tumor TMA cores, guided by an expert pathologist, are typically produced from a columnar tumor sample (or biopsy). Virtual columnar cores were taken at each chosen coordinate, and virtual TMA (vTMAs) were generated along these columnar cores. For each core, we quantified the error in PDAC composition between the first section of the core and any other given section (Fig 4B). We found that the relative error across 50 virtual cores increases dramatically within distances as low as 1.2 mm (240 tissue 4-μm sections) (Fig 4C). The mean relative error in cancer composition increases beyond 15% after 150 sections. We note that composition of tissue cores changes rapidly after merely tens of histological sections. This analysis shows the rapid decorrelation of PDAC content, even on expert-guided simulated biopsies on tumor region.

**Fig. 4.**
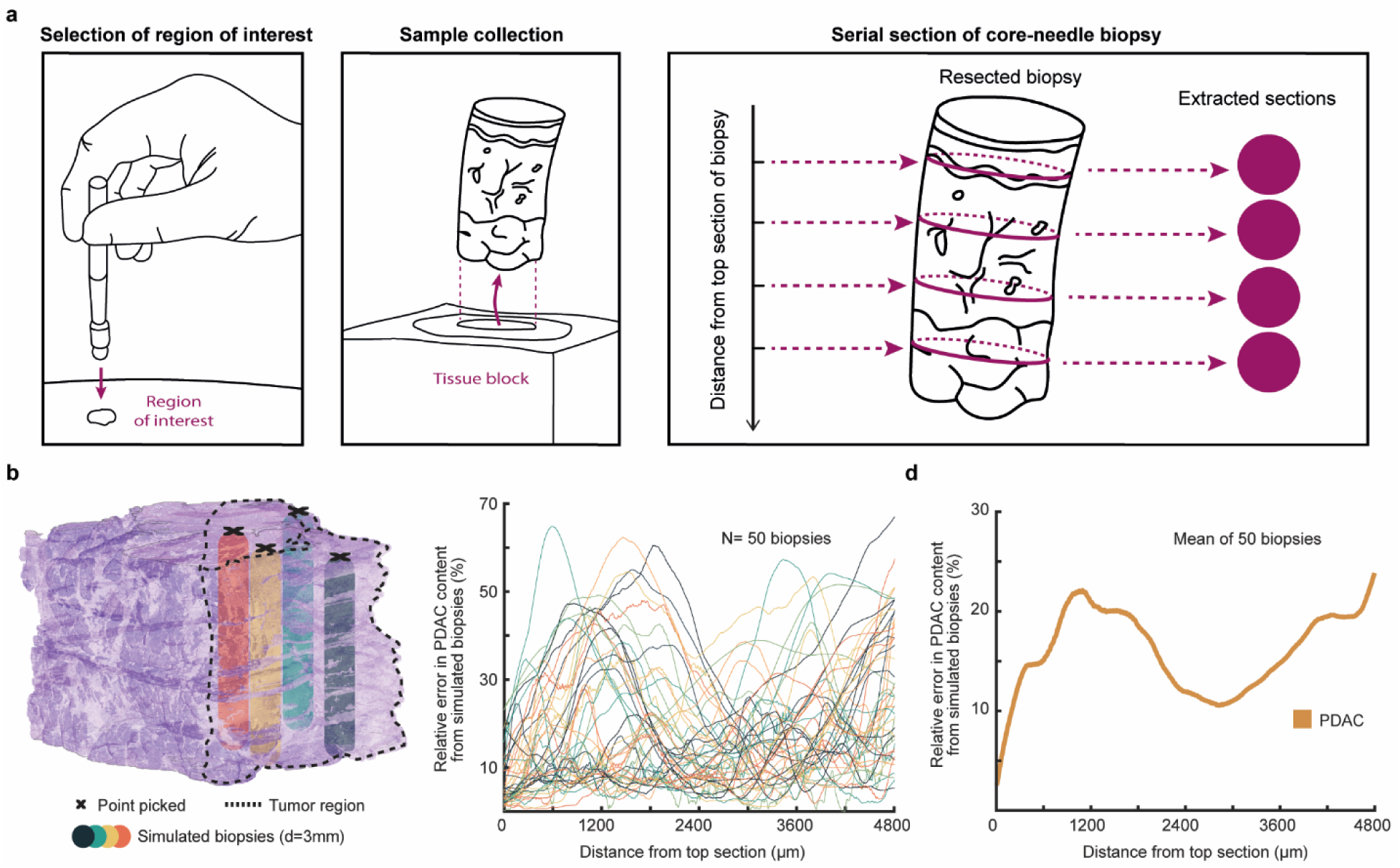
Error when TMA cores are used to assess spatial tissue content in a tumor. **(a)** Pathologist-guided selection of cores from virtual columnar cores (“biopsies”). **(b)** 50 target regions containing cancer were manually selected from the top slide for each of a block. **(c)** Virtual TMA (vTMAs) cores were obtained from the columnar cores, and the relative error in PDAC content between the top section and each vTMA was calculated. **(c)** Ensemble-averaged relative error in PDAC content for the 50 biopsies. The content in PDAC changes significantly along biopsies.

### Hundreds of TMAs are needed to capture the true tissue composition of WSIs and 3D tumors

We next aimed to understand the amount of information lost when subsampling a heterogeneous 3D tumor sample with a WSI or a TMA core of 1 mm in diameter instead of a full 3D assessment. To do this, we simulated virtual TMA (vTMAs) randomly located in the 2D WSIs and within the 3D-CODA samples (Fig 5A). We quantified the number of randomly sampled, non-overlapping vTMAs necessary to estimate the tissue composition of individual whole slide images within a preset error. For each simulation, between one and 100 vTMAs were generated, and the cumulative composition of eight microanatomical components of the pancreas (PDAC, fat, blood vessel, etc.) were compared to the overall content of these components of the WSI. This process was repeated for WSIs from all 64 individuals in the 2D-WSI cohort, and the resulting errors per simulation plotted as a function of the number of TMAs sampled. This process was also repeated to estimate the resulting errors per simulation as function of vTMAs within the seven 3D-CODA diseased volumes (Fig 5B), and the error when using randomly sampled, non-overlapping WSIs within the seven 3D-CODA volumes (Fig 5D).

**Fig. 5.**
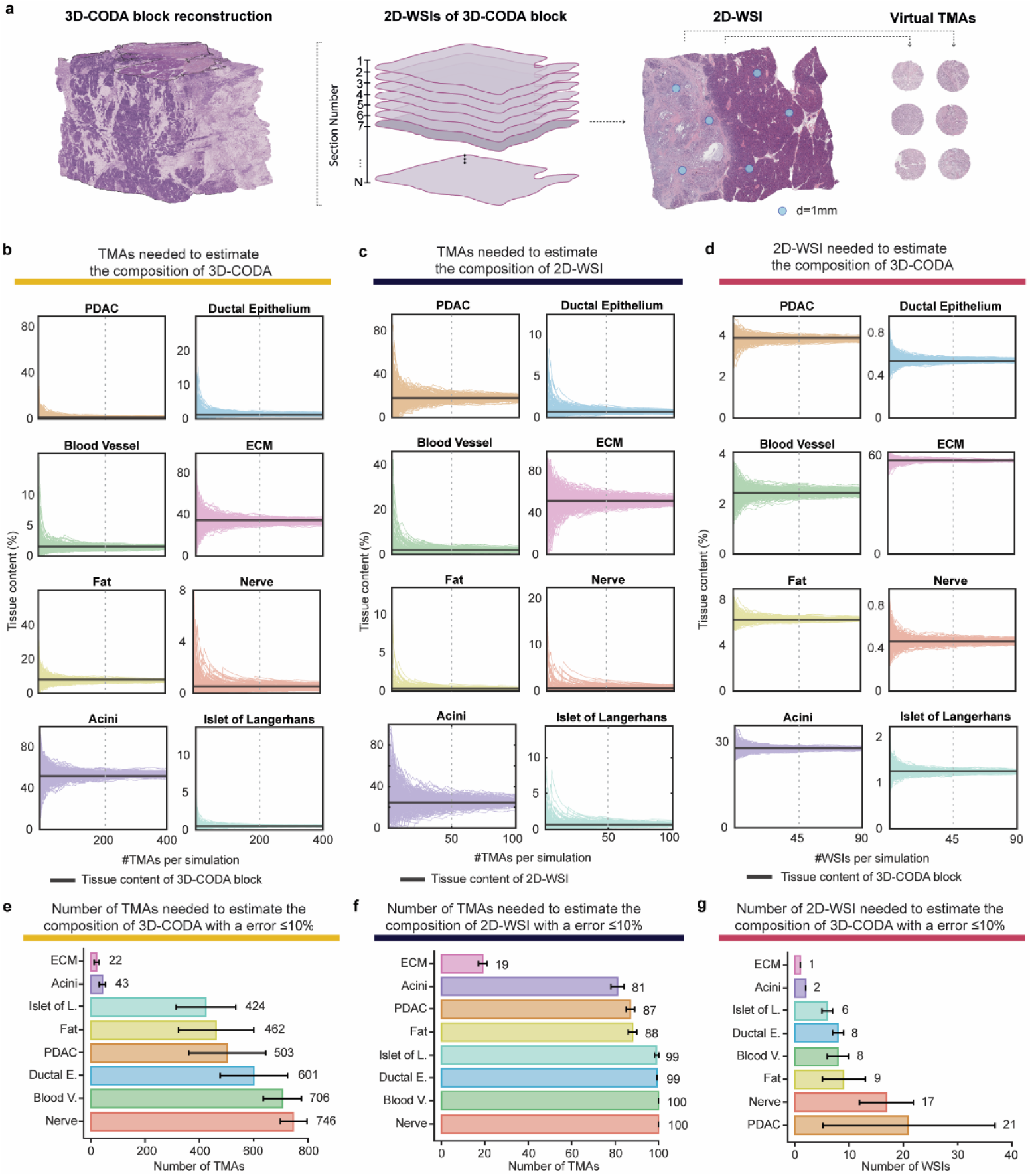
Hundreds of TMAs and tens of WSIs are needed to accurately estimate the true bulk tissue composition of 3D tumors. (**a**) Representation of 3D-CODA H&E-stained volume (left), 2D-WSIs in the 3D-CODA (middle), and virtual TMAs extracted from 2D-WSIs in the 3D-CODA. **(b)** The loss in the accuracy of calculation of tissue composition due to TMA subsampling was measured through 200 simulations of 1 to 100 virtual TMAs (vTMAs) in the 2D-WSI cohort. Tissue composition of the generated vTMAs was compared to the average 2D-WSI (black line). **(c)** The calculation of (a) was repeated for calculation of the loss in accuracy of estimation of tissue composition between TMAs (from 1 to 400 virtual TMAs) and 3D tumors. **(d)** The calculation of (a) was repeated for calculation of the loss in accuracy of estimation of tissue composition using actual WSIs and 3D tumors (of 1 to 90 actual WSIs). **(e, f, g)** Distilling the information from the simulations shown in (a-c), we calculated the number of TMAs and WSIs necessary to estimate WSI and 3D-tumor composition with ≤10% error. ECM showed the lowest error across all simulations, while sparser components, such as blood vessels, nerves, and PanIN required more (tens to hundreds) of tissue sections to reach ≤10% error.

For each simulation, increasing the number of TMAs for the rigorous assessment of bulk tissue composition in 2D images (Fig. 5C) and 3D samples (Fig. 5B) or WSIs for 3D samples (Fig. 5D), decreased the error of estimation of tissue composition, as expected. The number of simulated TMAs or WSIs necessary to reach pre-set error rates varied across different microanatomical tissue components of the pancreas. The number of TMAs or WSIs necessary to reach <10% error in estimation of true WSI or 3D-volume composition enabled the identification of low-heterogeneity and high-heterogeneity tissue components (Fig 5E, 5F, 5G). ECM consistently showed the lowest tissue heterogeneity, with an average of 19 TMAs necessary to reach <10% error in the estimation of 2D-WSI composition, an average 22 TMAs necessary to reach <10% error in the estimation of 3D-volume composition, and an average of 1 WSI necessary to reach <10% error in the estimation of 3D-volume composition. In contrast, accurate estimation of cancer composition required significantly more TMAs. The estimation of content in nerves revealed that these tissue components were the less prevalent, which required 100 TMAs for <10% error in estimation of 2D WSI composition, >700 TMAs for <10% error in estimation of 3D-volume composition.

This analysis showed how subsampling heterogeneous tumor samples with TMAs and WSIs can lead to information loss, when compared to whole >1cm^3^ 3D assessments. Simulations quantified the degree of dissimilarity between TMA and WSI sampling of each tissue component, when compared to the true bulk tissue content of the 3D samples.

### Quantification of spatial correlation and tissue composition variation at spatial transcriptomics scale

To characterize the spatial correlation decay across spatial transcriptomics region sizes, 65 total locations were selected across 7 tissue blocks. For each location selected, pixel-to-pixel correlation was computed from the initial WSI to progressively distant WSIs within the tissue blocks (Fig 6A). The spatial correlation for each individual region is shown in gray, and the mean of all computations is displayed as a colored line (Fig 6B). For tissue components including ducts, PDAC, islets of Langerhans, blood vessels, nerves, and fat, a decrease in spatial correlation of fifty percent was observed within just 40 microns (Fig 6C).

**Fig. 6.**
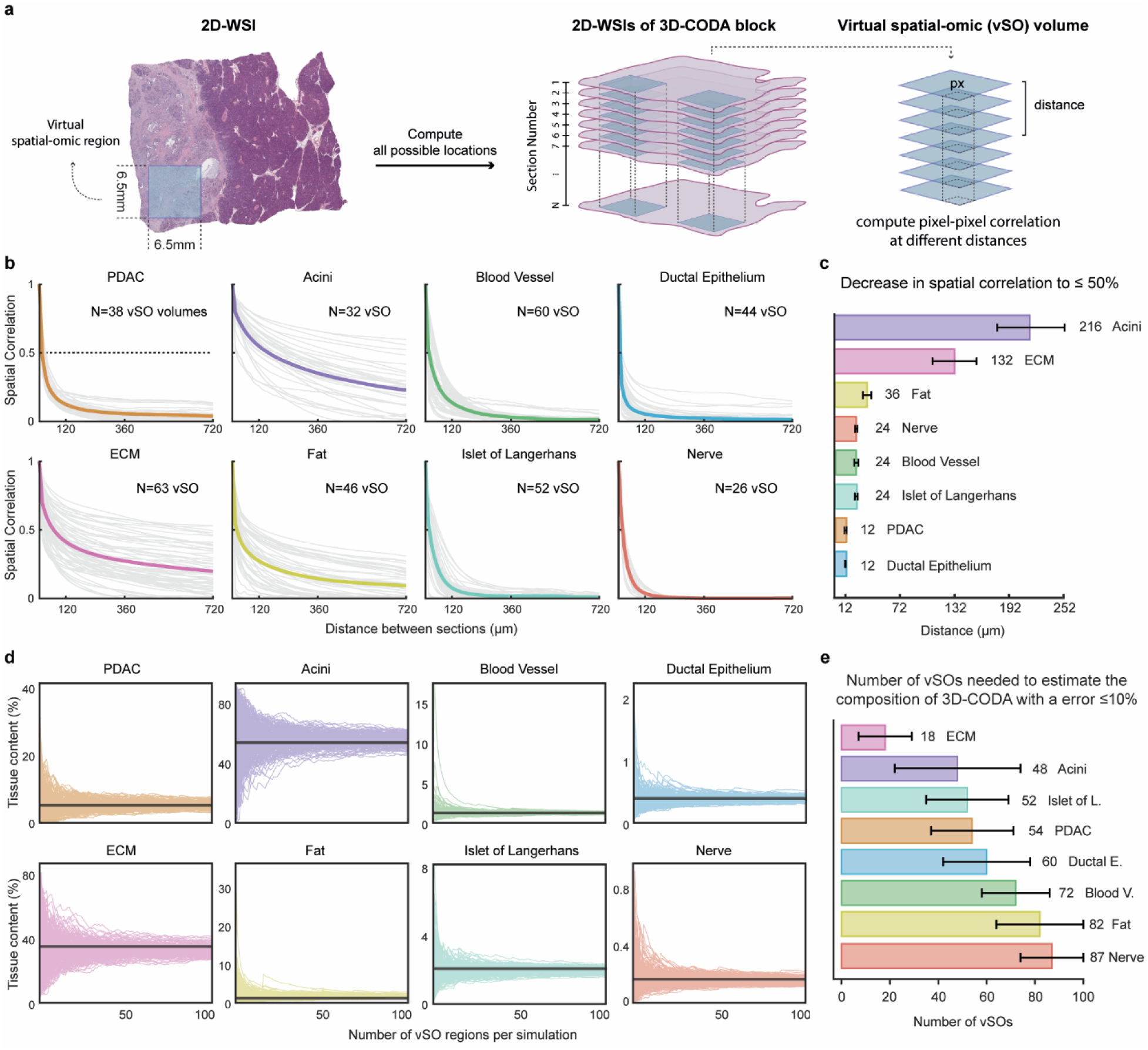
Quantification of spatial correlation and tissue composition variation at spatial transcriptomics scale. **(a)** 65 total locations (6.5 x 6.5 mm^2^) were selected across 7 tissue blocks. For each location, pixel-to-pixel correlation in tissue content was computed for all combination of pairs of spatial-omics size regions in that given 3D-CODA block. **(b)** The spatial correlation for each individual region is shown in gray, and the mean of all computations is displayed in color. **(c)** For tissue components including ducts, PDAC, islets of Langerhans, blood vessels, nerves, and fat, a decrease in spatial correlation of fifty percent was observed within just 40 microns. **(d)** The loss in the accuracy of tissue composition due to spatial-omic region subsampling was measured through 200 simulations of 1 to 100 virtual spatial-omics (vSOs) in the 3D-CODA cohort. Tissue composition of the generated vSOs was compared to the average 3D-CODA (black line). **(e)** Between 48 and 72 regions of (6.5 x 6.5 mm^2^) were required to estimate the tissue composition of components including acini, islets of Langerhans, PDAC, ducts, and blood vessels to within 10% error.

To more comprehensively understand the rate of spatial transcriptomics region size needed for accurate profiling tissue composition in whole tissue blocks, similar tissue composition simulations were computed for spatial transcriptomics size regions of 6.5 mm x 6.5 mm. By generating 6.5 mm^2^ regions from a 3D tumor volume, we computed the number of regions needed to recapitulate the true bulk tissue composition (Fig 6C). Our analysis demonstrated that for tissue components including acini, islets of Langerhans, PDAC, ducts, blood vessels required between 48 and 72 regions of 6.5 mm x 6.5 mm to estimate their true tissue composition content with less than 10% error (Fig 6D).

Comparative evaluation with previous simulations (Fig 5E, 5G) highlighted the superior capability of 2D-WSI to more comprehensively profile tissue composition from tissue blocks, followed by spatial transcriptomics regions, and lastly TMAs having the most limited sampling ability.

### Required sampling in 3D samples depends on the relative prevalence of the target tissue

Here, we assess the relationship between the sampling in a 3D tissue necessary to reach a preset error in the estimation of tumor (PDAC) content and the relative contents in invasive cancer and cancer precursor lesions PanIN in that tissue. For this calculation, we first utilized a previously reported cohort of 48 large 3D reconstructed samples of human pancreas tissue containing PanINs.^5^ We defined PanIN content as the volume percent of PanIN within the pancreatic ductal system: P_content_ = volume of PanIN / (volume of PanIN + normal ductal epithelium). Next, we calculated P_content_ for all possible combinations of consecutive slides subsampled from the above 3D cohort and calculated the relative error of the subsampled region to the P_content_ of the full 3D sample (Fig 7A). Visualizing this as bar plots for low, medium, and high P_content_ (0< X ≤33%, 33%< X ≤66%, and 66%< X ≤100%, respectively) revealed that few slides are needed to accurately determine the neoplastic content of samples containing large fractions of PanIN, while many slides are needed to accurately determine the neoplastic content of samples containing low fractions of PanIN (Fig 7B).

**Fig. 7.**
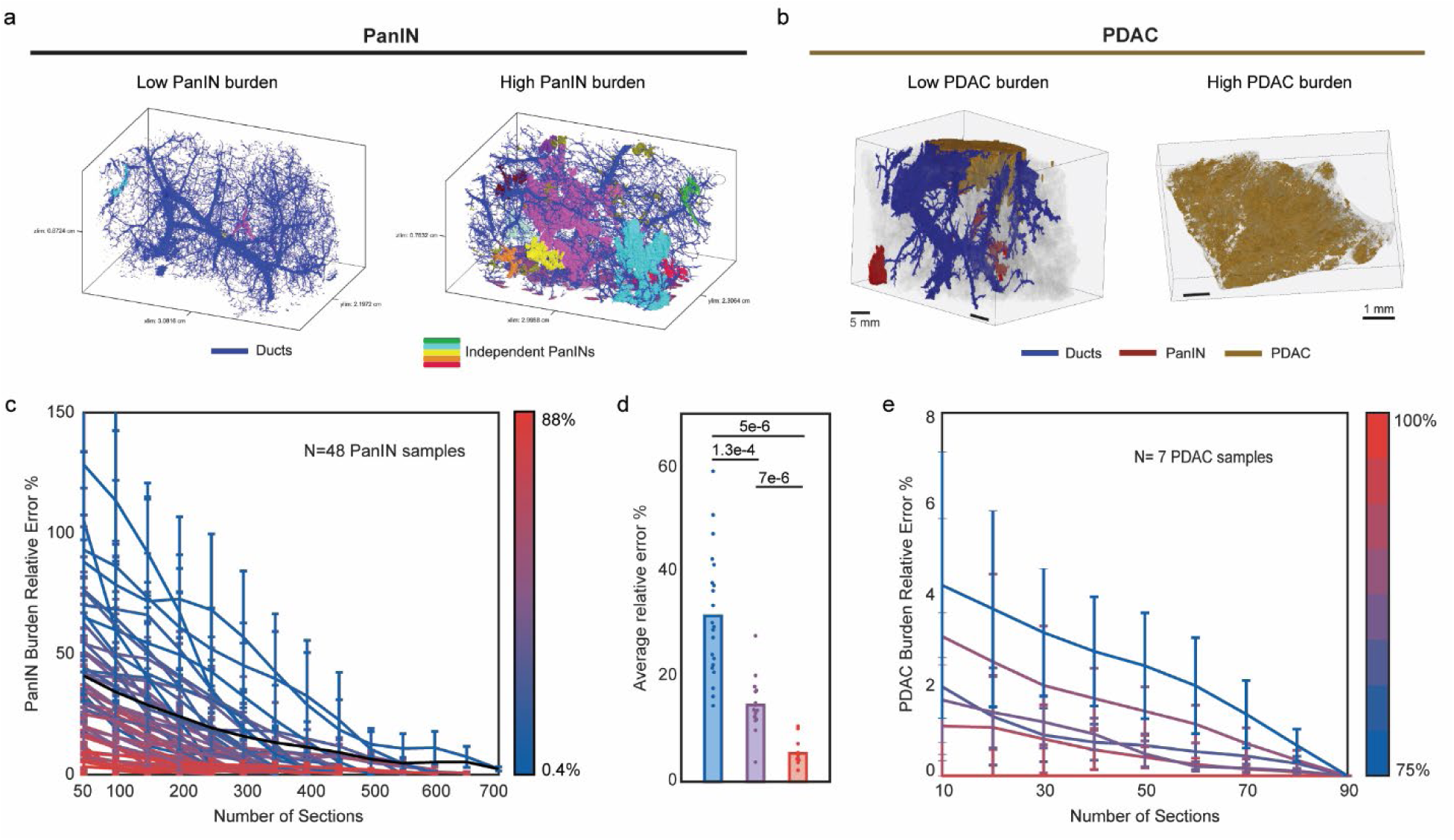
The number-of-WSIs-needed to capture the true bulk incidence in pancreatic neoplasms depends on their frequency in the sample. (**a and b**) 3D rendering of pancreata that contain low and high PanIN content (**a**) and low and high PDAC content (**b**). (**c**) For 48 3D pancreata containing PanIN (precursors to pancreatic cancer), the error in estimation of overall PanIN content as a function of the number of consecutive sections subsampled was computed. Lines are color-coded according to the overall PanIN content of the sample, from low (blue) to medium (purple) to high (red). (**d**) The data in (c) binned according to overall PanIN content to show that fewer sections are needed to accurately estimate the PanIN content of samples that contain many PanIN lesions, and *vice versa*. (**e**) The calculation of (c) was repeated for the seven samples containing PDAC in the 3D-CODA cohort to show that fewer sections are needed to accurately estimate cancer content in samples that contain high cancer composition, and *vice versa*.

After validating the PanIN content calculation on the cohort of 48 adjacent healthy tissue blocks, the same calculation was computed for the PDAC content on the initial seven 3D-CODA blocks obtained from individuals diagnosed with pancreatic cancer. We defined an analogous cancer content: C_content_ = volume of cancer / (volume of cancer + volume of PanIN + volume of normal ductal epithelium). Again, we found that few slides are needed to estimate the composition of cancer in samples with high cancer burden, but that many slides are necessary to estimate cancer composition in samples with low neoplastic content (Fig 7C).

These results suggest the rather intuitive guideline that the rarer the tissue component (e.g. PDAC content in samples from patients with low PDAC burden) to be studied is, the larger the number of WSIs is required for a rigorous assessment of that component content.

## DISCUSSION

Methods for spatially resolved cellular profiling have enabled in-depth quantitative mapping of tissues and tumors to study inter-patient and intra-patient differences in normal human anatomy and disease onset and progression. These methods profile extremely limited regions, which may impact the evaluation of tissue content and local heterogeneity due to tissue sub-sampling. Here, we apply CODA, a deep learning-based tissue mapping platform, to reconstruct the 3D microanatomy of surgically resected human pancreas specimens. To compare differences in the inter-and intra-tumoral heterogeneity in tissue content, we assess the spatially-resolved composition of a cohort of two-dimensional (2D) whole slide images (WSIs), and a cohort of 3D serially sectioned and reconstructed tissues of pancreata. We demonstrate the value of 3D by analyzing information loss and sampling issues when using subsampled 2D tissue sections. The spatial correlation in microanatomical tissue content decays significantly within a span of just a few microns within tumors. As a corollary, hundreds of TMAs, and tens of WSIs are required to estimate bulk tumor composition with <10% error in any given pancreatic tumor. The large error in the estimation of the occurrence of rare tissue components, for example the PDAC content in a sample with low PDAC neoplastic content, requires large numbers of WSIs to rigorously detect and measure these tissue components.

While 2D assessments using WSIs are typically sufficient for clinical diagnostic, they are insufficient for accurate assessment of spatially resolved tissue composition of organs and diseased tissues. A consequence is that digital pathology done on whole slides, which presumes that intra-organ variations in cellular content are smaller than patient-to-patient organ variations, may lead to erroneous conclusions, even in organs that are believed to be highly homogenous. In sum, we demonstrate that 3D assessments are necessary to accurately assess tissue composition and tumor content and provide guidelines for the rate of sampling necessary to rigorously assess spatially resolved tissue composition and associated tissue density and intercellular distances.

## Code and data availability statement

The data analyzed here is available from the corresponding author upon request. The code used to generate the 3D tissue maps is available on the following GitHub: https://github.com/ashleylk/CODA.

## Author contributions

A.L.K. and D.W. conceived the project. A.L.K. A.M.B., K.F., T.C., S.H., and R.H.H. collected and processed the human pancreas samples. A.L.K., A.F., E.V, S. J, V.M.R, A.J., and P.W. conducted the image analysis and heterogeneity quantifications. S.M, L.W, and R.H.H. validated histological analysis as clinical experts in pancreatic cancer pathology. A.F., A.L.K., R.H.H. and D.W. wrote the first draft of the manuscript, which all authors edited and approved.

---

**Fig. S1.**
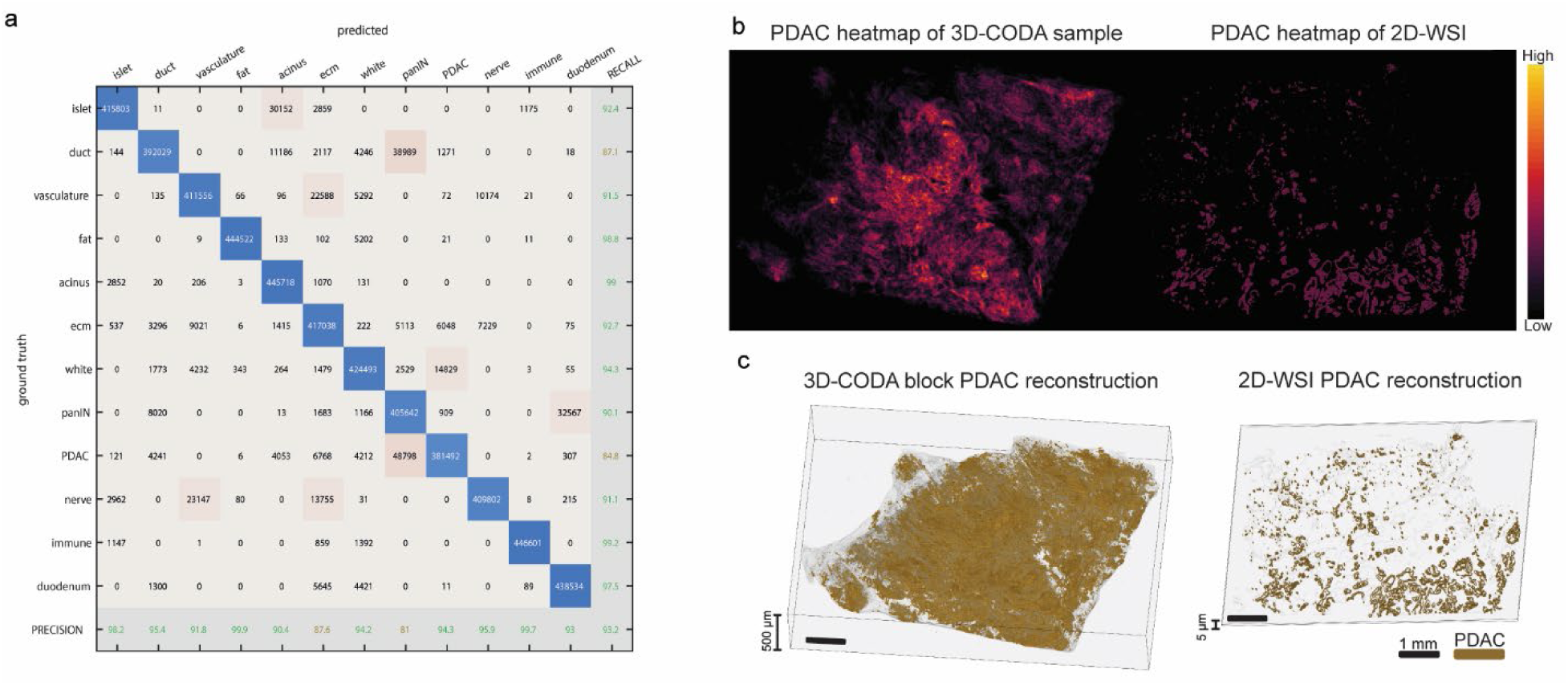
Validation of deep learning segmentation of pancreatic microarchitecture. **(a)** Confusion matrix representing the overall accuracy and per-class precision and recall of the segmentation algorithm used to map the tissue composition in the 2D and 3D samples. Confusion matrix built using independent manual annotations not used in training. **(b)** Z-projection of PDAC content in 3D volume as opposed to PDAC content in a single WSI. **(c)** 3D reconstruction of PDAC content in 3D-CODA block in comparison with single 2D-WSI.

**Fig. S2.**
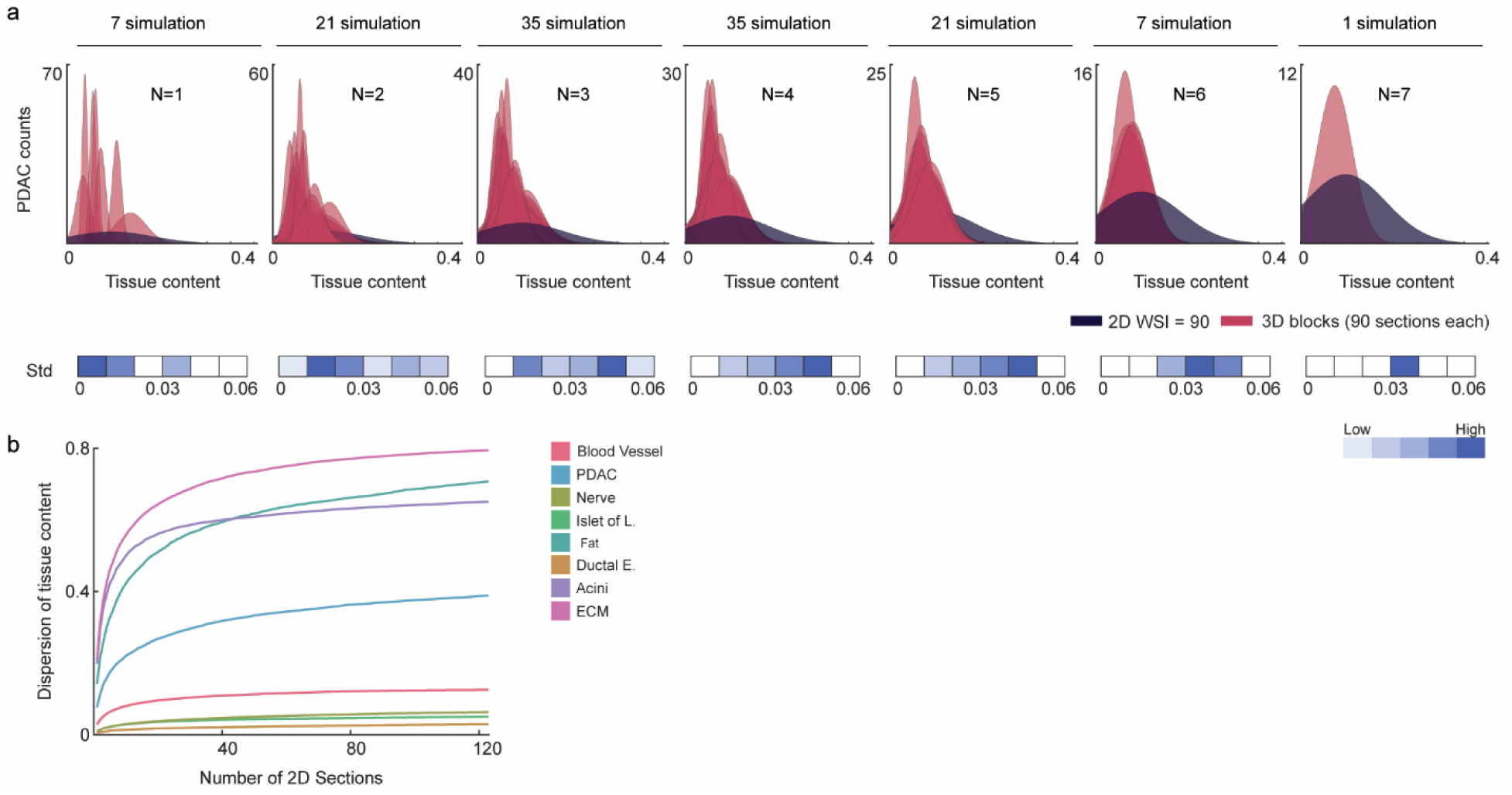
Number of patients necessary to capture the full range in tissue composition of PDACs. (**a**) Distributions in PDAC content for all combinations of individual 3D samples (red) compared to the full range in PDAC composition shown by the 64 patients (blue) in the 2D-WSI cohort. N is the number of 3D samples used in each plot. (**b**) Number of 2D WSIs (i.e. number of patients represented by one slide) necessary to saturate in inter-patient compositional heterogeneity. Composition curves saturate at ∼40-50 sections, justifying the number of 2D slides used for the 2D WSI cohort.

